# Chidamide combined with doxorubicin leads to synergistic anti-cancer effect and induces autophagy through inhibiting the PI3K/Akt/mTOR pathway in anaplastic thyroid carcinoma

**DOI:** 10.1101/2020.05.19.105288

**Authors:** Yishan He, Xinguang Qiu

**Affiliations:** Department of Breast Surgery, the First Affiliated Hospital of Zhengzhou University, Zhengzhou, China; Department of Thyroid Surgery, the First Affiliated Hospital of Zhengzhou University, Zhengzhou, China

**Author notes:** **Corresponding author:** Xinguang Qiu. No. 1 Jianshe East Road, Zhengzhou 450052, Henan, PR China.

**Keywords:** Anaplastic thyroid carcinoma, Chidamide, Doxorubicin, anti-cancer effect, autophagy

## Abstract

Anaplastic thyroid carcinoma (ATC) is a fatal malignant tumor, which belongs to the thyroid cancer with an overwhelmingly poor prognosis and eagerly demands effective systemic treatment strategies. We aimed to investigate the antitumor characteristics of Chidamide (CS055) in combination with doxorubicin (Dox) on ATC, and explore the underlying molecular mechanism. Herein, we found that CS055 and Dox inhibited proliferation, invasion and migration and promoted apoptosis of ATC cells. CS055 and Dox induced autophagic cell death (ADC) of ATC cell lines. And the expression of autophagy markers, BECN1, Atg5 and LC3-II was significantly enhanced in ATC cell lines treated with CS055 and Dox. Similarly, the *in vivo* study showed that CS055 and Dox administration significantly reduced tumor growth and induced tumor cell autophagy. Interestingly, the synergistic anti-cancer effect of CS055 in combination with doxorubicin was observed *in vitro* and *in vivo*. In addition, CS055 and Dox suppressed the proteins expression of p-P13K, p-AKT and mTOR *in vitro* and *in vivo* and combination of CS055 and Dox exhibited greatest inhibitory effect. Taken together, our findings concluded that CS055 in combination with Dox exerted antitumor activities and triggered autophogy in thyroid carcinoma probably through inhibiting the P13K/AKT/m/TOR signaling pathway.

## Introduction

Anaplastic thyroid carcinoma (ATC) is the a extremely biologically invasive neoplasm of malignant thyroid tumors with low incidence, rapidly disease progresses and low survival rate (Nagaiah et al., 2011). Because of its unstable genetic characteristics and complex genetic changes, the effective treatments have been lacking. Conventional methods such as surgical resection, radiotherapy and chemotherapy has no curative effect and cannot significantly improve the survival rate of patients, and it is particularly important to seek new treatment methods (Hallden and Portella 2012). At present, the biological treatment methods such as gene therapy, differentiation therapy, and protein inhibitor therapy are already in the exploratory stage (Fountzilas et al., 2017, Liu et al., 2018). Feasible induction of chemotherapy and radiotherapy sensitization often is applied in the comprehensive treatment therapy. The most commonly used chemotherapy drug for enhance the sensitization of chemoradiotherapy is doxorubicin (Dox), which was the only drug approved for anaplastic thyroid cancer treatment (Marano et al., 2017). Unfortunately, the valid time of doxorubicin efficacy is short, due to the resisting force to doxorubicin is rapidly formed once the treatment began (Wang et al., 2017). And the adverse effects of various chemotherapy drugs have resulted in insignificant clinical treatment, and the prognosis is poor. Therefore, the search for alternative drugs or drugs that enhance the sensitivity of chemotherapy drugs and reduce side effects is the direction for the future research.◻

HDAC inhibitor (HDACi) can selectively inhibit the activity of Histone deacetylase (HDAC) through binds to the catalytic region of HDAC, enhance the acetylation level of histones, loosen the chromatin condensation and restore the transcriptional activity. HDACi can blocks cell cycle, induce apoptosis, regulate autophagy, alter non-coding RNA expression, anti-angiogenesis, immune response, and effects on a variety of other cell signaling pathways to antitumor by altering epigenetic modification (Eckschlager et al., 2017). With the application of numerous HDACIs, its advantages of low toxicity, safety and broad antineoplastic spectrum have gradually been reflected, and it has become a major focus for the treatment of cancer.

Chidamide (CS055/ HBI-8000), a novel benzamide chemical type of HDACi, is currently being tested in many clinical trials in foreign and domestic country, which has been authorized to treat the cutaneous T-cell lymphoma and peripheral T-cell lymphoma in China (Gong et al., 2012). Studies have demonstrated that CS055 can lead solid tumors to apoptosis such as colon cancer (Liu et al., 2016), pancreatic cancer (Zhao and He 2015), non-small cell lung cancer (Zhou et al., 2014), hepatocellular carcinoma (Wang et al., 2012), even in other hematological tumors such as multiple myeloma (Xu et al., 2015), B-cell lymphoma (Lee and Kim 2012), leukemia cell line and primary cells (Gong *et al.* 2012) and leukemia stem cells (Li et al., 2015) have been certificated to promote apoptosis and induce cell cycle arrest. However, the effects and the mechanism of CS055 in anaplastic thyroid carcinoma have not been studied.

At present, drug combination therapy has become a hopeful therapeutic method for cancer treatment, specifically to drug resistant, and the synergistic effect of the two drugs helps to amplify weaker cell signals. Further, many studies have demonstrated that HDACIs could work in synergy to increase the antitumor activities of traditional chemotherapy drugs, of which included Vorinostat (SAHA), MS-275, Trichostatin A (TSA), and valproic acid (VPA) (Xie et al., 2010, Sung et al., 2011). The study of Zhang et al. (Zhang et al., 2017) has proved that there was a synergistic antitumor effect when used CS055 and Dox in combination with low dose concentration in peripheral T-cell lymphoma (PTCL). However, there is not relevant reports for the research of CS055 combined with Dox in ATC.

In this study, we explored the anti-tumor effect and possible mechanism of CS055, Dox or the combination of the two drugs in ATC cell lines and *in vivo* mice xenograft models. Our studies revealed anti-proliferation, anti-invasion, anti-migration and pro-apoptosis and inducing autophagy of CS055 and its synergistic effect with Dox, which might be involved in the P13K/AKT/mTOR pathway in ATC. Our findings provide a practical and theoretical basis for finding more effective treatments for anaplastic thyroid carcinoma.

## Results

### CS055 in combination with Dox synergistically inhibited cell proliferation and promoted apoptosis in ATC cell lines

Cell proliferation is one of the most striking features of Tumor cells activation. To test the effect of CS055 and Dox applied individually or in combination on ATC cell proliferation, FRO and 8505C were treated CS055 (1 μM) or/and Dox (0.5 μM) and then MTT assays were performed. Results showed that CS055 and Dox significantly restrained the proliferative activity of FRO and 8505C cells, and combination of CS055 and Dox had the strongest inhibitory effect (Figure 1A). Furthermore, as shown in Figure 1B, cell apoptosis analysis indicated that CS055 and Dox significantly induced cell apoptotic whether used respectively or in combination contrasted with the control. Combined treatment of CS055 and Dox resulted in highest percentage of apoptotic cells. These data suggest that combination of CS055 and Dox synergistically inhibits the proliferation and promotes apoptosis in ATC cells.

**Figure 1.**
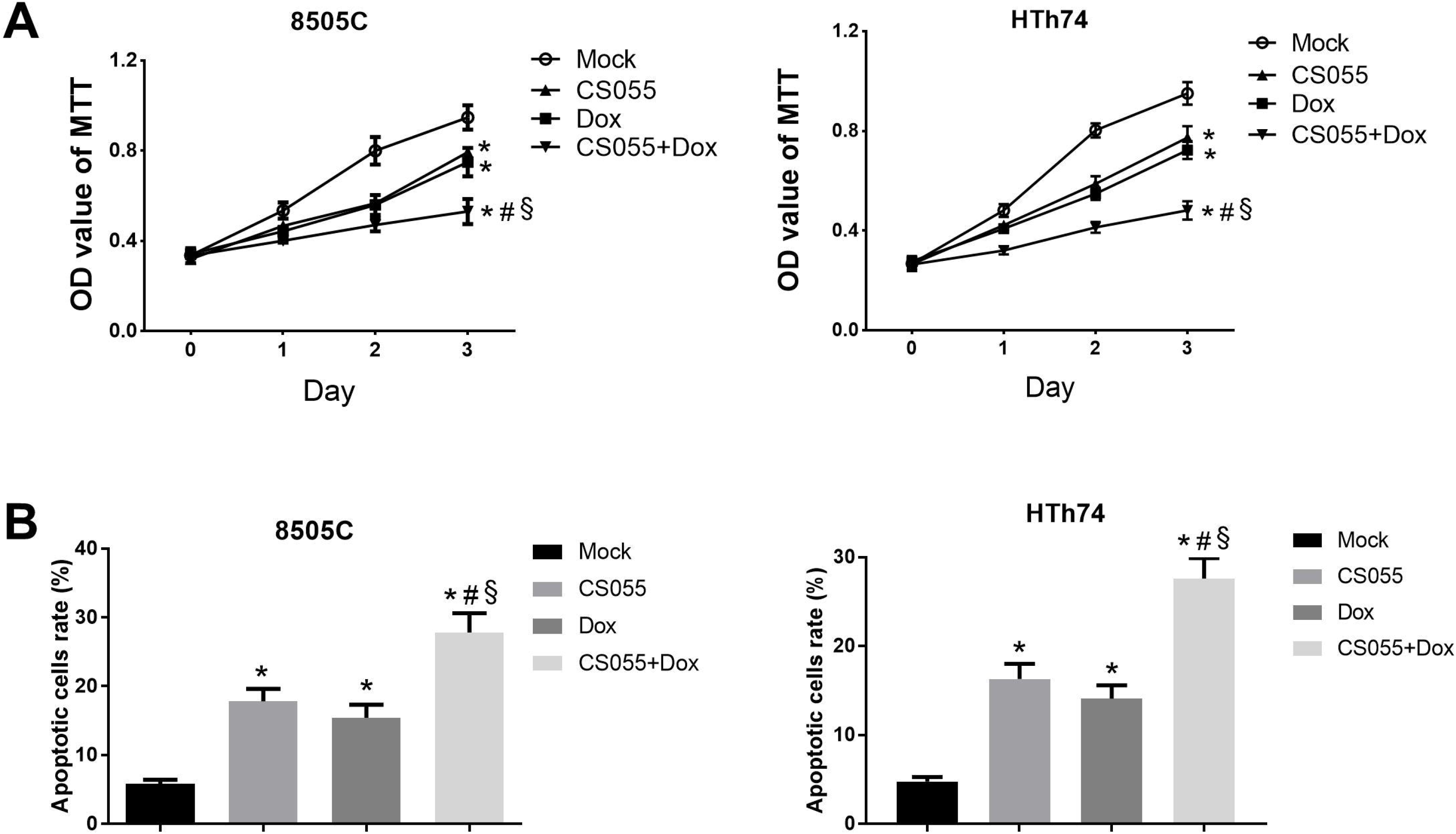
The effects of CS055 and Dox applied individually or in combination on cell proliferation and apoptosis of FRO and 8505C cells. A. FRO and 8505C cells were treated with CS055 (1 μM) and Dox (0.5 μM) for indicated times (0, 1, 2, 3 days) and the proliferative activity was measured by MTT assays. B. The treated FRO and 8505C cells were subjected to Annexin-V/PI staining with CS055 and Dox alone or in combination at indicated concentrations at 48 h and cells apoptosis was determined by flow cytometry. \**P* < 0.05 *vs.* the control group; ^#^ *P* < 0.05 *vs* CS055-treated group; ^§^*P* < 0.05 *vs.* Dox-treated group.

### CS055 in combination with Dox synergistically suppressed invasion and migration of ATC cell lines

Next, we probed into the effect of CS055 and Dox on the invasion and migration of ATC cell lines. As shown in Figure 2A, CS055 and Dox significantly suppressed the invasion of FRO and 8505C cells whether used alone or in combination. Moreover, CS055 and Dox presented a plus capacity of inhibition on invasion of FRO and 8505C cells. The results from wound healing assay showed that the treatments of CS055 and Dox significantly decreased FRO and 8505C cell migration compared to the control group. And compared with the single drug effects, the combination group significantly suppressed the cells migration capacity (Figure 2B). These outcomes demonstrated that combination of CS055 and Dox can synergistically inhibit the invasion and migration abilities of ATC cells.

**Figure 2.**
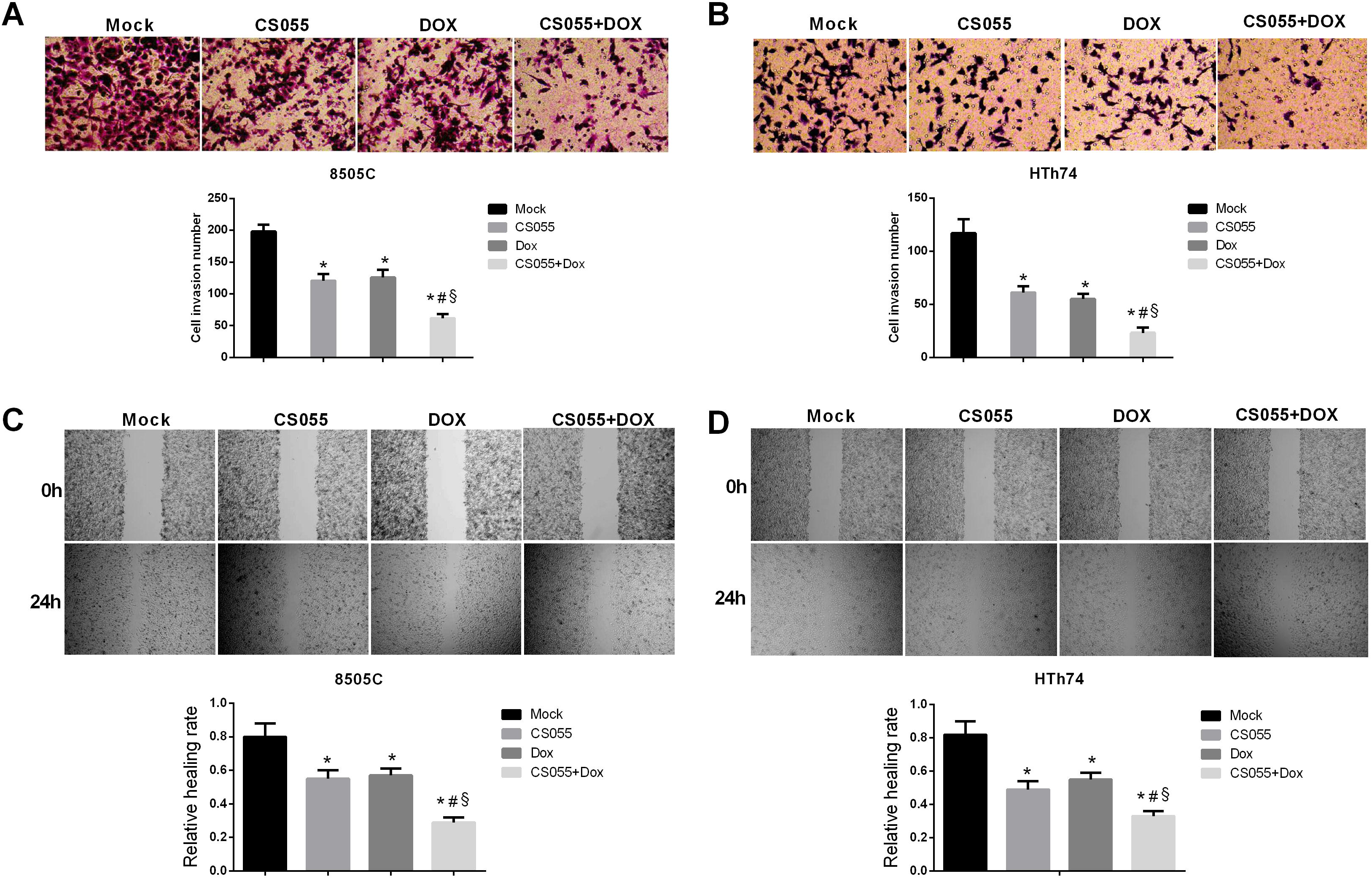
Cell invasion and migration were induced by CS055 and Dox applied individually or in combination in FRO and 8505C cell lines at 48 h. A. The cells invasion ability was determined with Transwells assay. B. The cells migration capacity was detected by wound healing assay at 0 and 48 h. \**P* < 0.05 *vs.* the control group; ^#^ *P* < 0.05 *vs* CS055-treated group; ^§^*P* < 0.05 *vs.* Dox-treated group.

### CS055 in combination with Dox synergistically induced autophagy in ATC cells

Autophagy, as another programmed cell death mode besides apoptosis, plays a significant role in maintaining the homeostasis of intracellular environment. As shown in Figure 3A, MDC staining (autophagic vacuoles) demonstrated that autolysosomes were highly accumulated in FRO and 8505C cells exposed to CS055 or Dox. The morphological characteristics demonstrated that autophagy was activated in FRO and 8505C cells since tested with CS055 and Dox. Moreover, CS055 and Dox showed an additive effect on MDC recruitment to autophagosomes in the cytoplasm of FRO and 8505C cells. To further verify the autophagy in ATC cells was induced by CS055 and Dox. The expression of BECN1, Atg5, LC3-I, and LC3-II proteins was detected by qRT-PCR and western blot assays. As shown in Figure 3B and 3C, indicated that the mRNA and protein expression levels of autophagy-related gene BECN1 (Atg6) and Atg5 were significantly up-regulated by CS055 and Dox treatment whether used alone or in combination and CS055 and Dox showed an additive effect. The transformation of the dissolvable form of LC3-I to the autophagic vesicle-associated form LC3-II is considered as a particular symbol of autophagosome formation. Results showed that CS055 and Dox significantly improved the proportion of LC3-II to LC3-I and CS055 and Dox showed an additive effect. These results indicate that CS055 works in synergy with Dox to activate autophagy of ATC cells.

**Figure 3.**
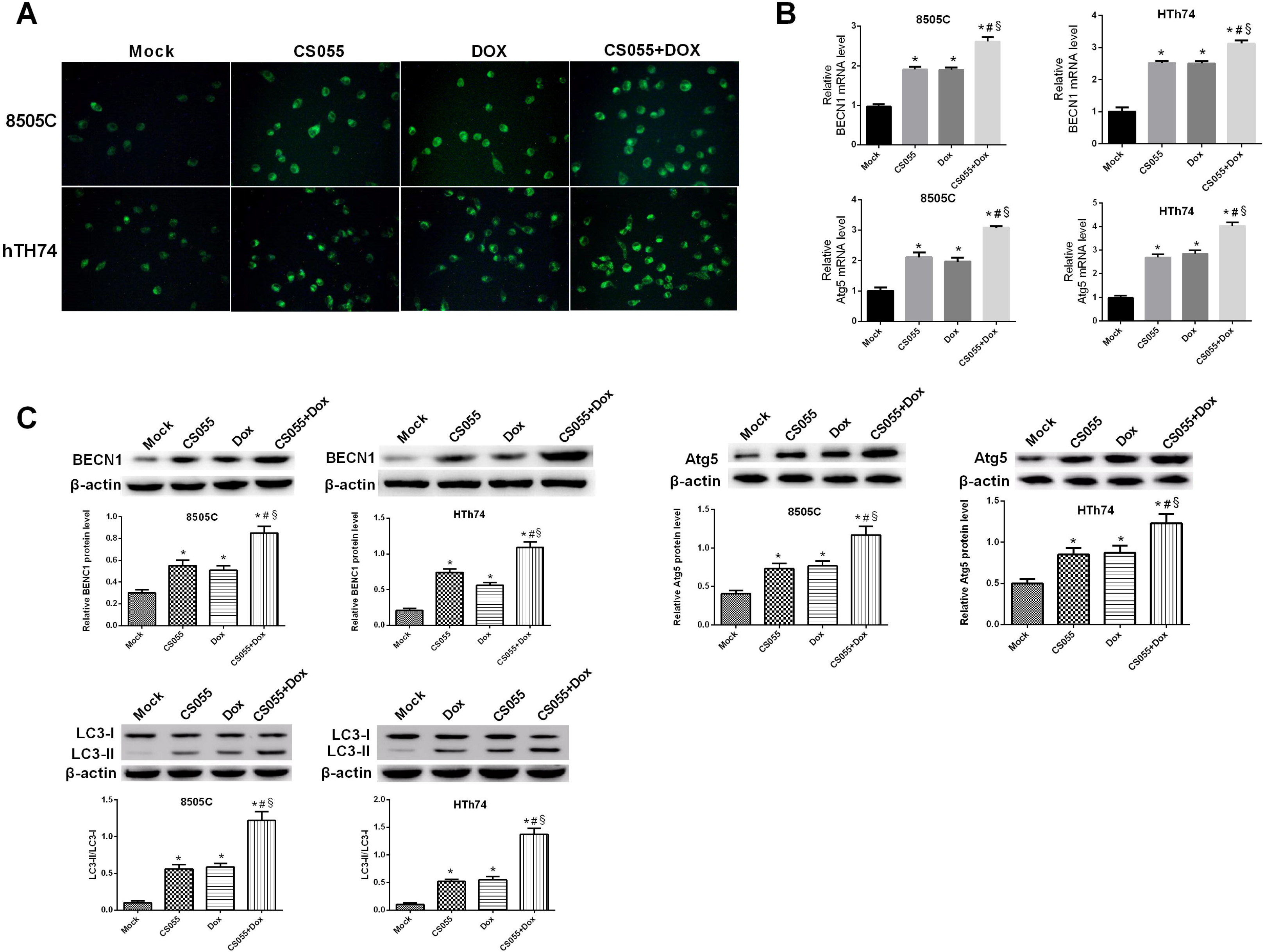
Cell autophagy induced by single CS055 or in combination with Dox in FRO and 8505C cell lines. A. Visualization of intracellular autophagic vacuoles in the FRO and 8505C cells after treatment with CS055, Dox, or CS055 and Dox in combination by MDC staining. FRO and 8505C cells after treatment with CS055, Dox, or CS055 and Dox in combination. Magnification ×200. B. The quantitative analysis of autophagy-related genes BECN1 and Atg5 expression in the FRO and 8505C cells by qPCR analysis. C. The expression of autophagy-related proteins BECN1, Atg5 and LC3-I/ LC3-II by western blot analysis. \**P* < 0.05 *vs.* the control group; ^#^ *P* < 0.05 *vs* CS055-treated group; ^§^*P* < 0.05 *vs.* Dox-treated group.

### CS055 in combination with Dox synergistically inhibited 8505C xenograft growth and induced autophagy in vivo

To validate the impact of CS055 and Dox *in vivo*, anaplastic thyroid carcinoma xenograft models were established by hypodermically injection of 8505C cells in nude mice. Figure 4A and 4B showed that CS055 and Dox significantly exerted inhibition of tumor growth, respectively. And CS055 in combination with Dox had significantly inhibitory effect on growth of xenograft compared to single drug effects. Furthermore, qRT-PCR and western blot analyses showed that CS055 and Dox significantly increase the proportion of LC3-II to LC3-I and enhanced the expression of BECN1 and ATG5 in 8505C cells xenograft tissues, and CS055 in combination with Dox had synergistical effect (Figure 4C and 4D). These results suggest that CS055 in combination with Dox synergistically inhibits ATC xenograft growth and induces autophagy *in vivo*.

**Figure 4.**
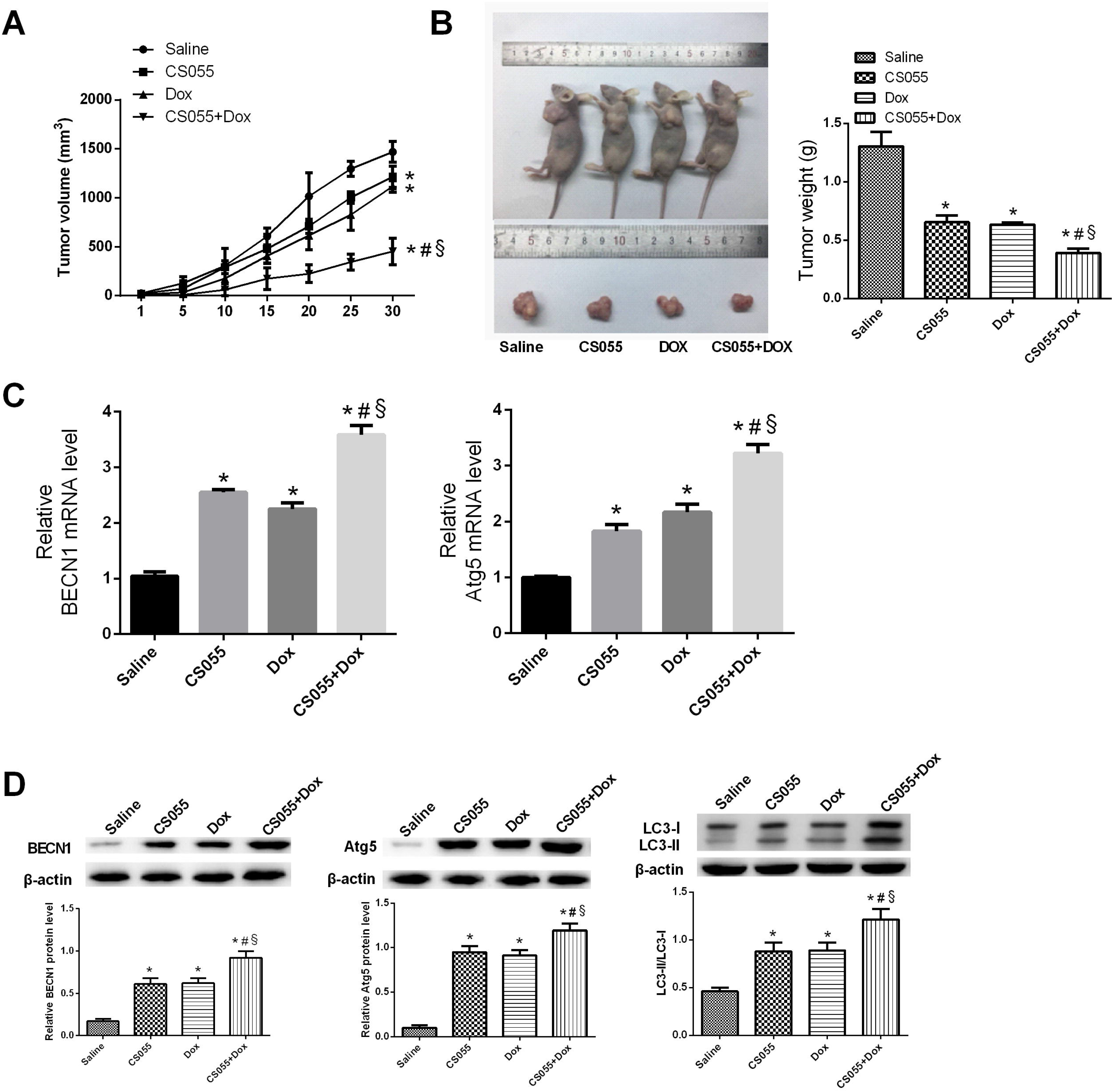
Effect of single CS055 and combination with Dox on the growth of ATC xenograft growth. A. Tumor volumes after CS055 (12.5 mg/kg/day) and Dox (3 mg/kg/day) alone or in combination treatment were indicated. B. Tumor weight was calculated. C. The quantitative investigation of BECN1 and Atg5 expression in tumor tissues of nude mice by qRT-PCR analysis. D. The protein expression of BECN1, Atg5 and LC3-I/ LC3-II in tumor tissues of nude mice by western blot. \**P* < 0.05 *vs.* the control group; ^#^ *P* < 0.05 *vs* CS055-treated group; ^§^*P* < 0.05 *vs.* Dox-treated group.

### CS055 in combination with Dox synergistically inhibited the P13K/AKT/mTOR pathway in vitro and in vivo

The P13K/AKT/mTOR signaling pathway is related to the administration of multiple cellular functions, containing proliferation, migration, cell growth, and autophagy (Sinclair et al., 2013, Abeyrathna and Su 2015). Western blot was applied to examine whether the P13K/AKT/mTOR signalling pathway was connected with the antitumor impact of CS055 and Dox on ATC cells and the xenograft models. As shown in Figure 5A and 5B, the protein expression levels of p-P13K, p-AKT and mTOR were significantly down-regulated after CS055 and Dox treatment whether used alone or in combination. Furthermore, there was a statistically significant in the combination therapy group compared with the single drug treatment group, which indicated that CS055 works in synergy with Dox to inhibit the P13K/AKT/mTOR signaling pathway. These data conclude that CS055 combination with Dox exerted antitumor activities and triggered autophogy in ATC probably through inhibiting the P13K-AKT-m-TOR signaling pathway.

**Figure 5.**
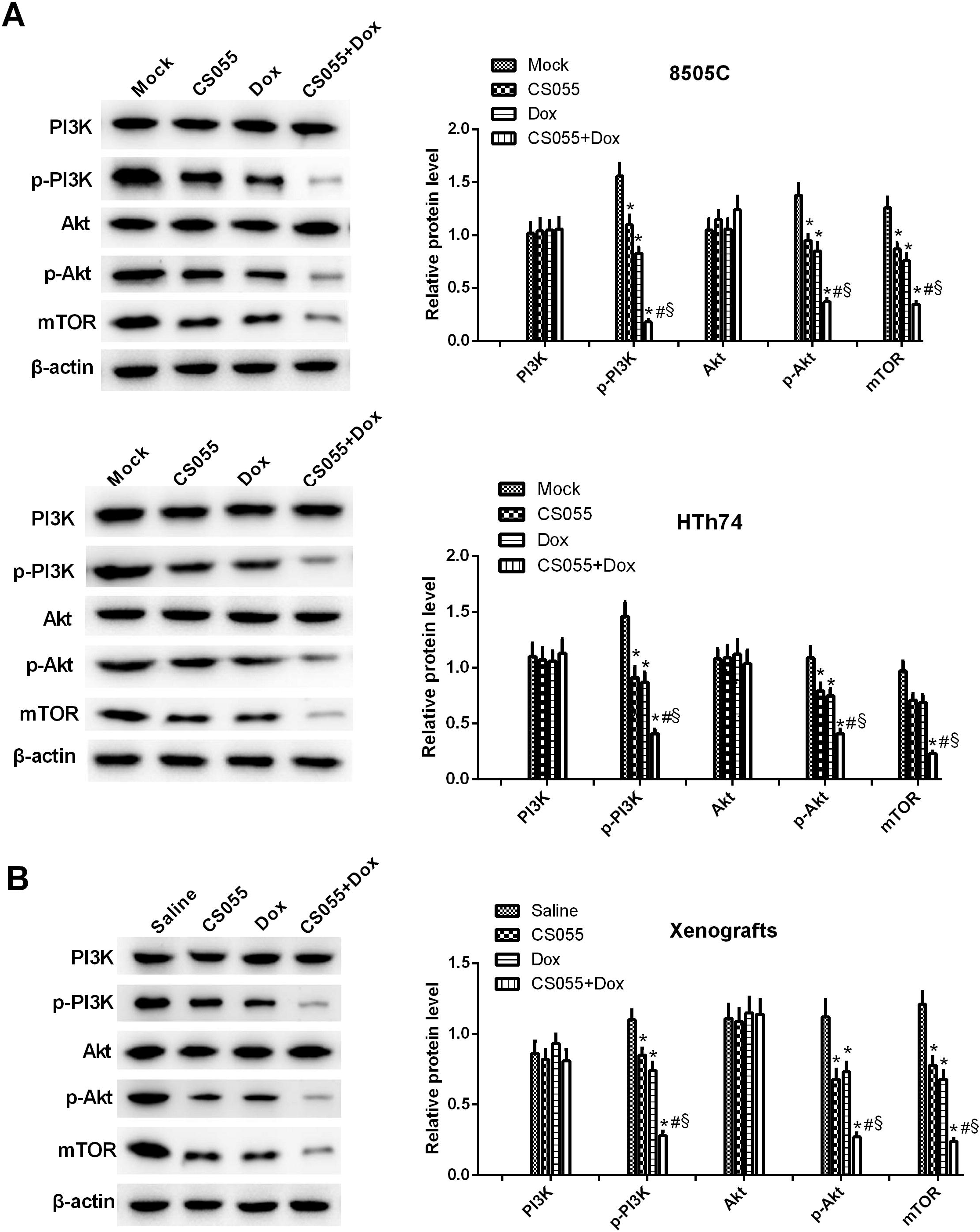
Effect of single CS055 and combination with Dox on P13K/AKT/m/TOR signaling pathway *in vitro* and *in vivo*. A and B. The protein expression of related P13K/AKT/m-TOR pathway was detected by western blot in FRO and 8505C cells and xenograft tissues with CS055 and Dox treatment whether used alone or in combination. \**P* < 0.05 *vs.* the control group; ^#^ *P* < 0.05 *vs* CS055-treated group; ^§^*P* < 0.05 *vs.* Dox-treated group.

## Discussion

Previous studies given evidence of the impacts of HDAC inhibitors in miscellaneous tumor cells *in vitro* and *in vivo*, which implied that HDACi is a promising new tactic for the therapy of cancers (Cappellacci et al., 2018). It has been widely shown that the anti-cancer effects of HDACi are mainly obtained through the following ways: inducing the generation of reactive oxygen species (ROS); inducing the growth arrest, differentiation and apoptosis of tumor cells; preventing angiogenesis and lymphangiogenesis; and preventing the invasion and migration of tumors (Suliman et al., 2012). Since then, the US FDA approved the clinical application of SAHA, which belongs to HDAC inhibitor, for CTCL therapy. This field of study then gets even hotter and inspires us to seek and design more available and effective HDAC inhibitors.

CS055, a histone deacetylase inhibitor developed and synthesized independently in China, is mainly targeted on subtypes 1, 2, 3 of class I HDAC and subtypes10 of class IIb HDAC, which can inhibit the cell cycle and induce cell apoptosis of tumors through epigenetic regulation mechanisms (Gong *et al.* 2012, Yao et al., 2013), restore the drug sensitivity of drug-resistant tumor cells and inhibit the metastasis and recurrence of tumors (Zhou *et al.* 2014, Gao et al., 2017). Studies have shown that when CS055 combined with gemcitabine acts on pancreatic cancer cells, it can synergistically enhance gemcitabine-induced cell growth arrest and apoptosis, and enhance the anti-tumor effect of gemcitabine (Qiao et al., 2013). In our study, we exploited the combination treatment of CS055 and Dox to test ATC cells for purpose of exploring the anti-tumor effect and potential mechanisms. Our results showed that CS055 and Dox induced apoptosis and inhibited proliferation in ATC cells. Moreover, the experimental results demonstrated that CS055 and Dox used in combination synergistically enhanced the antitumor effects on ATC cells.

The degradation of extracellular matrix (ECM) is the primary condition for breaking the barrier between inter-tissues and facilitating the migration of tumor cells across tissues. HDACi is reported to prevent the metastasis and infiltration of cancer cells by regulating the transcription and expression of metastasis-suppressing genes and metastasis-promoting genes (Pratap et al., 2010, Koshkina et al., 2011). Studies reported by Lin et al. have showed that CS055 restrains the migration of lung cancer cells in a dose-dependent manner (Lin et al., 2016). Lu et al (Lu et al., 2018) found that combination of CS055 and Ganoderma *in vitro* inhibits proliferation, invasion and infiltration abilities of A375.S2 cells and induces a large number of apoptotic cells of A375.S2 cells with a dose dependent manner. Consistent with that, our results demonstrated that CS055 and Dox significantly suppressed the invasion and migration of FRO and 8505C cells and worked in synergy.

Autophagy, as an important mechanism for maintaining homeostasis of the internal environment, plays a significant role in the pathogenesis, progress, therapeutic strategies and metabolism of tumors, and can improve the sensitivity of thyroid cancer cells to chemoradiotherapy (Morani et al., 2014). A study has shown that the multiple tumor cell death caused by suberoylanilide hydroxamic acid (SAHA), a HDACi, is independent of caspase-activated apoptosis, but induces autophagic cell death through a large number of activated autophagy reactions, thus inhibiting the growth of tumor cells (Lee et al., 2012). HDACi can increase the formation of intracellular acidic vesicles, recruited LC3-II to autophagosomes, promote the expression of Beclin1 protein and decrease the expression of p62 (Gandesiri et al., 2012). Zhao et al (Zhao and He 2015) have reported that the autophagy of Pa Tu8988 pancreatic cancer cells significantly increased after the treatment of CS055, which suggesting that CS055 can promote pancreatic tumor cell death by inducing autophagy. In this study, CS055 and Dox significantly repressed the growth of the tumor whether used alone or in combination. Moreover, CS055 and Dox induced autophagic vacuoles and increased the expression levels of BECN1, Atg5 and ratio of LC3-II to LC3-◻*in vitro* and *in vivo* study and there was a synergistically effect in the combination treatment group compared with the single drug group, which confirmed that the autophagic process seems to be another probably mechanism to induce cancer cell death treated with CS055 and Dox in ATC tumor.

Studies have shown that P13K/AKT/m/TOR signaling pathway is closely related to the occurrence and development of human tumors, which plays a significant role in malignant proliferation, metastasis, apoptosis, invasion and autophagy and other functions of tumor cells (Guan et al., 2007, Zhang et al., 2011, Xie et al., 2013). Therefore, PI3K-Akt-mTOR signaling pathway is expected to become a new target for the treatment of malignant tumors. Study presented by Tam-burrino et al. (Tamburrino et al., 2012) has demonstrated that the PI3K/Akt/mTOR signaling pathway was over-activated in the occurrence and progression of thyroid cancer. Therefore, targeting this pathway can effectively inhibit the proliferation of ATC cell lines. Moreover, HDACi can down-regulating the P13K/AKT/m/TOR signaling pathway and inducing apoptosis and autophagy (Chiao et al., 2013, Foster et al., 2014). Encouraged from the previous studies mentioned above, we tested the expression of proteins involved in P13K/AKT/m/TOR signaling pathway in ATC cells or xenograft nude mice. As indicated in our western blot results, the protein expression levels of p-P13K, p-AKT and mTOR were significantly down-regulated after CS055 and Dox treatment whether used alone or in combination, and more significantly, the decrease in the combination treatment group was statistically significant compared with that in the single drug groups, suggesting that CS055 and Dox inhibit the ATC tumor probably through inhibiting the P13K/AKT/m/TOR signaling pathways.

## Conclusion

In conclusion, *in vitro* and *in vivo* experiments shown that the combination treatment with CS055 and doxorubicin can synergistic inhibited the proliferation, invasion and migration and induced apoptosis and autophagy of ATC cells, all of which probably through inhibiting the P13K/AKT/m/TOR signaling pathways. These suggest that CS055 combination with doxorubicin may be a promising clinical therapeutic regimen for the anaplastic thyroid carcinoma, which might shed new light on anaplastic thyroid carcinoma therapy.

## Materials and methods

### Cell lines and cell culture

The human anaplastic thyroid carcinoma cell lines FRO and 8505C were afforded by Biological Cell Bank of Shanghai Academy of Chinese Sciences. The cell lines were cultured in RPMI-1640 (GIBCO. Uxbridge, UK) comprised of 10% fetal bovine serum (FBS; GIBCO) and 1% penicillin /streptomycin (GIBCO). All cells were maintained in a humid atmosphere containing 5% CO_2_ at 37°C.

### Assessment of cell proliferative viability by MTT assay

Briefly, cells (2×10^3^ cells/well) were cultured in triplicate in 96-well plates in the presence or absence of CS055 (Shenzhen Chipscreen Biosciences Ltd, Shenzhen, China) and Dox (Sigma, St. Louis, MO, USA) with the final volume of 100 μl. After treatment for the indicated times (0, 1, 2, 3 days), 10 uL of MTT (Sigma) (5 mg/mL) was added to each well with that incubation time for 4 h at 37°C.Then we removed the supernatant discreetly, added 100 μL DMSO to each well, shaken the plate until the crystals were dissolved compliantly. Finally, the absorbance was evaluated in Varioskan™ LUX Multi-function microplate reader (Thermo Scientific, USA) at a wavelength of 490 nm. The reported data derived from three separate experiments.

### Detection of apoptosis by flow cytometry analysis

In the study, we used the An Annexin-V-fluorescein isothiocyanate (FITC) Apoptosis Detection Kit (BD Biosciences, USA) to detect the cells apoptosis using flow cytometry. Cells were seeded with the density of 2 × 10^5^ cells/well in six-well plates and treated with CS055 and Dox. After incubation for 48h, the cells were gathered and the cell suspension was coordinate to 1 × 10^6^ cell/ml. The apoptosis of FRO and 8505C cell lines was monitored via Annexin V-FITC/propidium iodide (PI) double staining. Finally, the treated cells were analyzed in a Beckman Gallios flow cytometer. All experiments were implemented in triplicate.

### Invasion and migration assays

Cell invasion assays were implemented in 24-well plates with Costar Transwell with 6.5 mm diameter and 5 mm pores (Corning Costar Corp., NY, USA). Cells were seeded in the superstratum chamber of Boyden compartments at a density of 5×10^4^ cells/well with serum free media containing CS055 and Dox individually or together. The substratum chamber was contained with 600μL of 0.2% BSA-supplemented RPMI-1640 medium for 24 h at 37 °C and 5% CO_2_, followed by immobilization and dyeing by the use of crystal violet (0.1%).The invaded cells were photographed.

Cell migration was detected by wound healing assay. Cells were clutured in the 6-well plates at 2 × 10^5^ cells/well in RPMI-1640 complemented with 10% FBS and grown to 75–85% confluency. The sterile micropipette spear was used to eradicate the single-deck cells and the plates were washed with PBS. Then, the culture solution was substituted by free-serum. The migration ability was detected and imaged after 0 or 24 h of incubation under the inverted microscope (Olympus IX51, Tokyo, Japan). The migration rate of ATC cells was analyzed using ImageJ software (National Institutes of Health, MD, USA).

### RNA extraction and Quantitative reverse-transcriptase polymerase chain reaction (qRT-PCR)

Total RNA of the ATC cell lines and mice tumor tissue was extracted according to the RNA^iso^TM Plus kit instruction manual, of which then used as the template for first-strand cDNA synthesis. qPCR was performed on a LightCycler 480 II (Roche Diagnostic Ltd., Basel, Switzerland) with RealStar Green Power Mixture (GenStar Biosolutions Co., Lt., Beijing, China) according to the manufacturers’ instructions. Gene expression level was computed by the following the equation: relative gene expression =2^−(treated(ΔGENE−ΔGAPDH)−control(ΔGENE−ΔGAPDH))^. *GAPDH* was used as the endogenous control to standardize mRNA content. Each swatch was experiment in triplicate.

### Western blot analysis

The treated ATC cells and nude mice tumor tissues were collected and dissociated with RIPA lysis buffer. The total protein lysates were extracted and protein concentrations were caculated by Bicinchoninic acid (BCA) assay. The equal protein specimens were separated by SDS-PAGE and then electro-transferred onto the nitrocellulose membrane. Membranes were blocked for 30 min and incubated with specific antibodies (against BECN1, Atg5, LC3-◻/LC3-◻, P13K, p-P13K, AKT, p-AKT, mTOR, and β-actin) overnight at 4’C following hatch with secondary antibodies for 2 h. Protein signals were imagined by ECL detection reagents according to manufacturer’s instruction using a ChemiDoc XRS system (Bio rad, Hercules, CA, USA). ALL specific antibodies were obtained from Cell Signaling Biotechnology (Danvers, MA, USA).

### MDC incorporation assay

Cells at a density of 2 × 10^5^ cells/well were cultured in a 6-well plate and incubated with the indicated drug treatment for 48 h. After washing with cold PBS, autophagic vacuoles were then stained with 0.05 mM monodansylcadaverine (MDC), which was an autofluorescent base with the capacity of backlogging in autophagic vacuoles, at indoor temperature for 30 min. After incubation, cells were imagined using a Laser confocal fluorescence microscope at 600X magnification (Olympus IX70, Tokyo, Japan).

### In vivo experiment

Forty BALB/c nude mice at the age of 5 weeks (18-22 g) were offered by the Animal Center of the Chinese Academy of Sciences. Experiments were approved by Animal Experimentation Ethics Committees of the First Affiliated Hospital of Zhengzhou University. 8505C cells (2 × 10^7^/200 μl/mouse) were suspended in PBS and hypodermically injected into the flank of the nude mouse to establish the anaplastic thyroid carcinoma mouse model. Mouse with approximately similar tumor size were stochastically separated into 4 treatment groups (n=10): control (vehicle), CS055 (12.5 mg/kg/day), Dox (3 mg/kg/day) and (CS055 12.5 mg/kg + Dox 3 mg/kg/d) groups and were given through intraperitoneal (ip) once a day consecutively for 30 days. Tumors were gauged every five days. The tumor volumes were measured and calculated with the following formula: volume (mm^3^) = (width ×width^2^)/2. After 30 days, the treated mice were euthanized and the tumors were collected and examined.

#### Statistical analysis

Data were analyzed using statistical software SPSS 18.0. All values were presented as the mean ± SD and the statistical analyses were implemented via the Student’s *t*-test. And p-values <0.05 was considered as statistically significant.

## Acknowledgements

No.

